# Neuronal basis of audio-tactile speech perception

**DOI:** 10.1101/2024.08.16.608369

**Authors:** Katarzyna Cieśla, Tomasz Wolak, Amir Amedi

## Abstract

Since childhood, we experience speech as a combination of audio and visual signals, with visual cues particularly beneficial in difficult auditory conditions. This study investigates an alternative multisensory context of speech, and namely audio-tactile, which could prove beneficial for rehabilitation in the hearing impaired population. We show improved understanding of distorted speech in background noise, when combined with low-frequency speech-extracted vibrotactile stimulation delivered on fingertips. The quick effect might be related to the fact that both auditory and tactile signals contain the same type of information. Changes in functional connectivity due to audio-tactile speech training are primarily observed in the visual system, including early visual regions, lateral occipital cortex, middle temporal motion area, and the extrastriate body area. These effects, despite lack of visual input during the task, possibly reflect automatic involvement of areas supporting lip-reading and spatial aspects of language, such as gesture observation, in difficult acoustic conditions. For audio-tactile integration we show increased connectivity of a sensorimotor hub representing the entire body, with the parietal system of motor planning based on multisensory inputs, along with several visual areas. After training, the sensorimotor connectivity increases with high-order and language-related frontal and temporal regions. Overall, the results suggest that the new audio-tactile speech task activates regions that partially overlap with the established brain network for audio-visual speech processing. This further indicates that neuronal plasticity related to perceptual learning is first built upon an existing structural and functional blueprint for connectivity. Further effects reflect task-specific behaviour related to body and spatial perception, as well as tactile signal processing. Possibly, a longer training regime is required to strengthen direct pathways between the auditory and sensorimotor brain regions during audio-tactile speech processing.

## Introduction

In most everyday situations we perceive speech as a multisensory signal, combining inputs from hearing and vision. Already from an early age newborns learn to associate speech sounds with visually available cues from lip-reading and gestures, and congruent audio-visual stimuli are preferred over the non-congruent ones already at several months of age (Hauswald et al. 2018; Hickok et al. 2018). A corresponding visual input facilitates understanding especially in challenging acoustic situations, such as in background noise and/or when the sound is distorted (e.g. Sumby & Pollack, 1954; Ross et al. 2011, Guediche et al. 2014. Crosse et al, 2016), This benefit is especially critical for the hearing impaired population, including users of cochlear implants (a neuronal prosthesis for sensorineural hearing loss), some of whom despite almost perfect hearing in quiet, find it extremely challenging to e.g. distinguish between multiple concurrent speakers (review in Bernstein et al. 2022).

The audio-visual speech advantage is believed to be driven by multi-stage integration, where temporal cues from visual inputs are first fed into the auditory cortex directly (potentially through direct connections between primary sensory regions), followed by further steps facilitating recognition of specific linguistic units (e.g., phonemes, syllables) and at the final stage disentangling the meaning (Bourguignon et al, 2020; Peelle and Sommers, 2015; O’Sullivan et al. 2021, Hauswald et al. 2018). The reported brain regions associated with integration of audio-visual speech (but not only) signals include the posterior superior temporal sulcus (pSTS), superior marginal gyrus/angular gyrus, inferior occipito-temporal cortex, anterior insula, middle temporal visual area (MT/MST), Extrastriate Body Area (EBA) and subregions of the frontal cortex, all corresponding to encoding different aspects of spoken communication (Beauchamp et al. 2004, Hertrich et al. 2020; Beer et al. 2013; Nath et al. 2012; Bernstein et al. 2008). Electroencephalographic (EEG) studies show that tracking auditory speech envelope in the auditory cortex also improves through concurrent lip-reading, which effect is most profound in noisy environments (e.g. Bourguignon et al, 2020; Crosse et al. 2016; O’Sullivan et al. 2021). In the current study we set out to explore another multisensory context of speech comprehension, and namely audio-tactile. Specifically, we provide distorted speech signals through hearing and additional low-frequency speech-extracted vibrations on the skin (<200Hz; Bolanowski et al. 1988). As opposed to the familiar audio-visual speech context, the audio-tactile pairing is utterly new to the participants and untrained either during the critical periods of development or during evolution (Heimler and Amedi, 2020).

Using touch to convey speech information has been a subject of investigation for quite some time, originally conceptualized as an aid for people with hearing deficits (before development of modern cochlear implants). Starting with the Tadoma method (Alcorn, 1932) in which a person places a palm over the mouth and jaw of the speaker to perceive vibrations, a number of dedicated electro- and vibrotactile aids were developed in 70s/80s transmitting various aspects of speech signals on the skin (e.g. Galvin et al. 2001; Cowan et al. 1989). When tactile vibrations were introduced, certain hearing-impaired individuals showed improved ability to differentiate speech features, particularly when supplemented with speech-reading cues. Several studies from our lab and others have also demonstrated that specifically designed tactile vibrations, extracted from a continuous speech signal, can improve understanding of sentences in noise, when delivered on the fingertips. This effect was found for both the typically hearing individuals, as well as among users of cochlear implants (Fletcher et al. 2018; Ciesla et al. 2019; Ciesla et al. 2022; Huang et al. 2017). In parallel, two recent EEG studies examined simple audio-tactile speech perception. Riecke and colleagues (2019) showed improved tracking of distorted auditory speech signals (but not improved intelligibility) when tactile speech-shaped envelopes were delivered on the skin as vibrations, while Guillemont and colleagues (2022) demonstrated that tactile pulses synchronized with the rhythm of syllables improve comprehension of words in noisy conditions.

Audio-tactile crossmodal interactions are observed extensively in behavioral studies for various simple stimuli (e.g. Crommet et al. 2019; Ro et al., 2009; Yau et al., 2009, Gori et al. 2014; Russo et al. 2019). The loci of multisensory integration of have been found in the auditory cortex, pSTS, anterior insula, as well as the parietal operculum (secondary somatosensory cortex), depending on the task and the manipulated features of the stimulus (e.g., Renier et al. 2009; Caetano and Jousmäki, 2006; Kassuba et al., 2013; Landelle et al. 2023; King et al. 2019; Foxe et al. 2009). There has been minimal research published on the neuronal correlates associated with the integration of audio-tactile speech inputs specifically or the brain’s encoding of tactile speech units in general. In probably the only fMRI study investigating this topic, it was shown that trained vibrotactile speech stimuli engage speech-related subregions in the superior temporal gyrus (STG), but only if the stimuli preserve the temporal dynamics of the applied auditory speech (Damera et al. 2023).

Here we perform a series of behavioral tests, as well as a functional magnetic resonance (fMRI) experiment during which typically hearing individuals are asked to repeat sentences. An MR-compatible device delivering tactile vibrations to the fingertips had been developed specifically for this study (Ciesla et al. 2019; Ciesla et al. 2022). To make the task more demanding, we used speech signals which are vocoded, i.e. distorted both in frequency and amplitude (as in the cochlear implant system), and also presented against speech-like noise. The goal is to pose a challenge resembling the one frequently faced by users of cochlear implants in everyday situations, i.e. understanding speech in presence of competing signals. Actual cochlear implant users cannot undergo fMRI examination, due to the electromagnetic interferences (ASHA 2007). At the same time we aimed to increase the probability of multisensory enhancement, based on the inverse effectiveness rule of multisensory integration, which posits that the benefit from the additional sensory information is proportional to the reliability/clarity of the input delivered through the main sensory modality (Meredith & Stein 1983; Holmes & Spence 2006; Crosse et al. 2016). In addition, we introduce a condition in which multisensory enhancement should diminish, by using vibrotactile stimuli that are non-congruent with the auditory speech signal (i.e. correspond to a different sentence). Between the two fMRI sessions, the subjects participate in a dedicated audio-tactile speech training session. Previous studies have shown that typically hearing people can indeed learn to understand distorted speech/speech in noise through training (e.g. Erb et al., 2013; Casserly and Pisoni, 2015), as well as learn to benefit from visual cues (here *tactile*) in acoustically challenging speech contexts (e.g. Bernstein et al. 2013).

We speculate, based on previous findings of our lab and by others, that the performance in the speech understanding task will improve when auditory speech is combined with congruent vibrotactile stimuli, as compared to audio only or when paired with a non-congruent tactile speech input. This would indicate successful integration, specifically after a dedicated training session. With respect to the underlying neuronal mechanisms, we hypothesize that audio-tactile speech integration should engage processes similar in nature to that of audio-visual integration. We propose that the process might involve basic synchronization of the two signals, both with respect to the onset-offset of the target sentence and potentially also its exact temporal and frequency fluctuations, with the temporal structure in the tactile signal further helping segment the auditory signal into meaningful components for better comprehension (O’Sullivan et al. 2021). In addition, with vibrations representing literally the same time-varying signal, the integration should potentially be rapid (Crommett et al. 2019). At the same time, however, even though audio-tactile interactions are abundant in everyday experience (e.g. during listening to music, or when feeling the vibration of a ringing mobile phone), the specific multisensory speech context is utterly novel. Therefore, we expect to see the following effects at the neuronal level : 1) first the participants will rely on an established familiar connectivity network for audio-visual speech integration (involving regions, such as e.g. MT, EBA, LOC) which after dedicated training might become “less visual” and “more tactile”, as the participants will rely more on the novel tactile inputs (Makin and Krakauer 2023); 2) training will result in enhancement of communication between early auditory and early sensory regions, directly or through an early integration region, such as pSTS (which also specializes in processing temporal signals, King et al. 2019); 3) for the (potentially) multi-level audio-tactile speech integration, we speculate to also see involvement of regions such as the parietal SMG/AG (attentional aspects of integration) and frontal cortices (decision making), with possibly weaker effects in the non-congruent vs congruent audio-tactile test condition (Kassuba et al. 2013).

## Material and Methods

All the study procedures were in accordance with the Declaration of Helsinki from 2013, and were approved by the Ethics Committee of the Hadassah Hospital (protocol no 353-18.1.08). Participants provided informed consents and were compensated for their participation. All data was collected at the premises of the ELSC Neuroimaging Unit in Jerusalem, Israel, with the MR data acquired in a 3T Skyra MRI scanner by Siemens and a 32-channel head-coil. The timeline of the experiment is shown as Table 1.

**Table 1.**
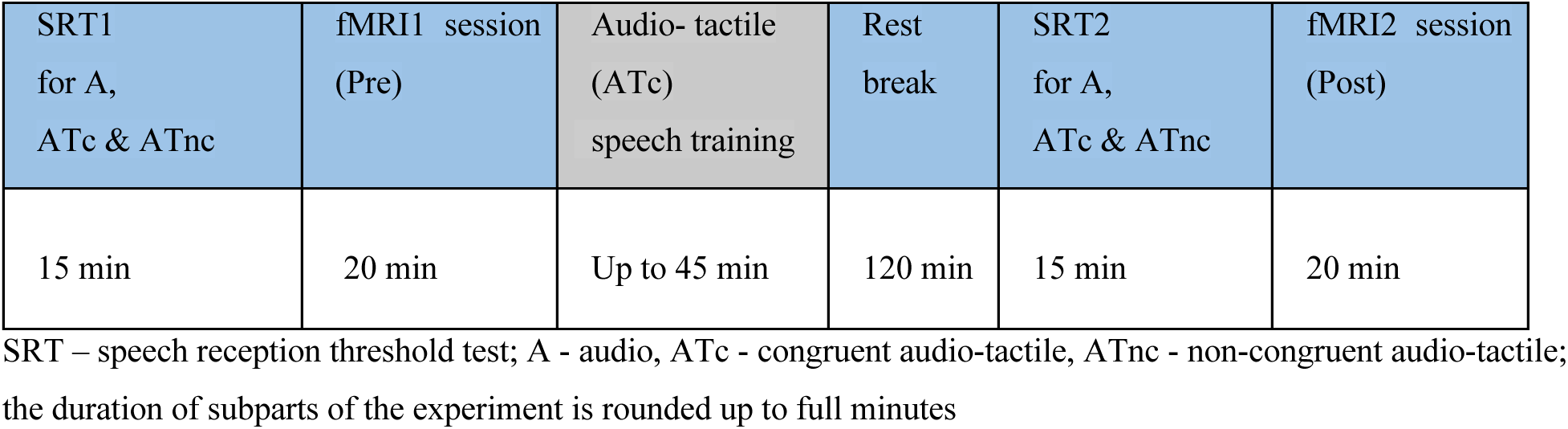
Timeline of the experiment.

### Participants

Twenty (20) typically hearing participants (6 male/14 female, 27.05+/-4.3 years, range 22-35) were recruited for the study. The participants were all right-handed and reported no history of neurological or neurodevelopmental impairments. MRI data collected in 3 participants had large image artifacts (due to the subject’s motion or signal drop-out), beyond repair, and were therefore excluded. The reported data comes from 17 individuals (6 male/11 female, 26.8+/-4.2).

### Experimental procedures

#### Stimuli and estimation of the Speech Reception Threshold

Before the first fMRI exam and following a short practice with the set-up, each participant had a speech-in-noise comprehension test, to estimate his/her individual Speech Reception Threshold (SRT, i.e. SNR between the target sentence and the background noise, for 50% understanding) in three comprehension tasks : a) auditory sentences (A), b) auditory sentences accompanied with congruent tactile vibrations on two fingertips (ATc), b) auditory sentences accompanied with non-congruent tactile vibrations on two fingertips (ATnc). The sentences were in English, spoken by a male, derived from the HINT database (Nilsson 1994). All sentences had a simple semantic content, such as e.g. “It’s getting cold in here” or “The chocolate cake was good”; each was 2.2-2.5s in duration. They were all presented against simultaneous background noise (International Female Fluctuating Masker, IFFM; Ehima 2016). In all participants, the first test was A, followed either by ATc or by ATnc (in half of participants), in both rounds i.e. before and after training. For each test, a different set of 20 sentences was used. The task of the participants was to repeat the sentences, one by one. The initial SNR for both SRT tests was set at 0 which corresponded to 65 dB SPL for both target and noise. Then an adaptive procedure was applied, increasing the SNR by 2 dB when the person was unable to repeat the whole sentence, and decreasing the SNR by 2 dB if the response was correct (Levitt, 1978). The response was considered correct only if the person repeated each word of the sentence exactly (except for personal pronouns, such as he/she that were both considered correct). The task finished after 20 sentences were played with no repetitions, and the final SNR was calculated as the mean of 5 last reversals. The sentences in the experiment were vocoded using a procedure developed at the World Hearing Center in Warsaw, Poland which entailed: bandpass filtering to 8 channels, signal rectification, low-pass filtering, envelope wave extraction (100–7500 Hz) with a 6th-order bandpass filter, narrowband noise modulation, simulating electrode interactions (through spread of excitation profiles), summation (Walkowiak et al., 2010). The energy of all sentences was then normalized with a standard RMS procedure and peaks of energy were leveled to –6 dB. For the congruent test condition, the tactile vibrations represented low frequencies of the current auditory sentence (ATc), while for the non-congruent test condition, the vibrations were extracted from another random sentence in the HINT database (ATnc). For vibrations, the fundamental frequency was first extracted from the sentence using the STRAIGHT algorithm (Kawahara et al., 1999). The amplitude information for the f0 (pitch) contour was extracted by low-passing the original signal with a 3rd-order bandpass digital elliptical filter with cut-off frequencies equal to the highest and the lowest frequency in the f0 contour, and then was normalized to its maximum digital value (0 dB attenuation) to provide maximum intensity of vibration across the whole f0 frequency range. This procedure resulted in the congruent and non-congruent sentences having different envelope content. For all test conditions the person heard the sentences through noise-canceling headphones (BOSE QC35 IIA) and felt vibrations on two fingertips (index and middle) of the right hand. The vibrations were delivered through an in-house developed device with two piezoelectric plates, one for each of the two fingertips (index and middle; see Fig 1; more details in : Ciesla et al. 2022). Before the test, it was made sure that each participant could feel the vibrations on their fingertips.

**Figure 1.**
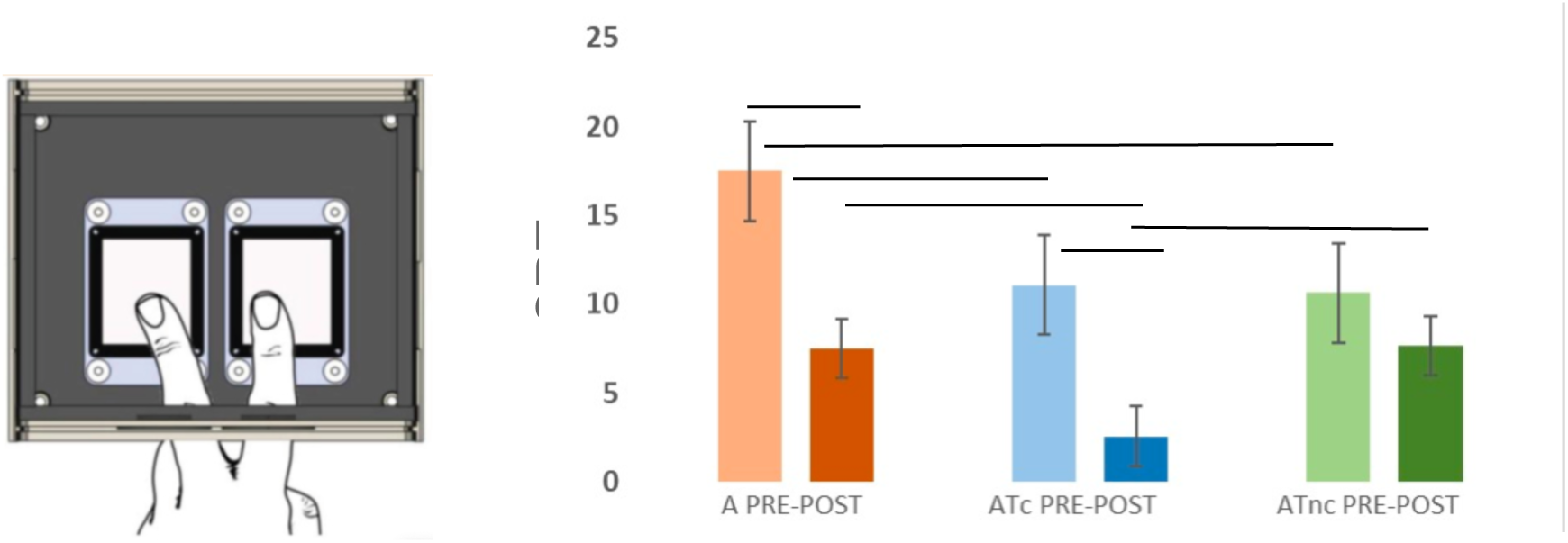
Mean results in the speech comprehension task before (PRE) and after (POST) Training; SNR - speech to noise ratio for 50% understanding; whiskers represent standard error of the mean; A - Audio, ATc - Audio-Tactile congruent, ATnc - Audio-Tactile non-congruent; on the left, a depiction of two fingertips on the piezoelectric plates of the vibrating device; *** p<0.001 unc., ** p<0.01 unc.

#### Speech comprehension task in MRI

The participants did the same task of speech comprehension inside the MR scanner, as they did outside of it. The sentences were vocoded and presented against background noise, at an SNR (SRT) determined individually for each person and each condition. There were six fMRI experimental runs, three before and three after training. As in the SRT tests, the order of conditions was swapped for half of the participants, between ATc and ATnc, following the A condition. In each fMRI run, i.e. audio before training (A PRE), audio-tactile congruent before training (ATc PRE), audio-tactile non-congruent before training (ATnc PRE), audio after training (A POST), audio-tactile congruent after training (ATc POST), audio-tactile non-congruent after training (ATnc POST), a different list of 24 sentences was used (predetermined), to avoid learning effects. Sentences were played, one by one, for approximately 3 seconds, followed by 3 seconds for the participant to repeat them. The sentences were presented through MR-compatible headphones and the participant responded to a pneumatic microphone attached to the head coil (OptoAcoustics). Three sentences in a row were followed by a rest baseline block of the same duration, i.e. approximately 22.5 s (3 x TA, see below). The researcher recorded the responses for further evaluation. There was no feedback given to the participants during the fMRI experiment. Participants were asked to keep their eyes focused on a cross on the screen. During a few-minute scanning trial at the very beginning, all confirmed that they can hear the sentences well and that they can feel the vibrations on their fingertips. The parameters of the task-fMRI scanning sequence were the following : SPARSE BOLD data acquisition, TR =7.5s, TA = 1.5s, TE=32.6ms, voxel size : 2×2×2 mm, 72 slices, flip angle 78, FOV 192 mm, Bandwidth 1736 Hz/voxel, iPAT off, no acceleration. One fMRI run was 6:38 min in duration, during which 48 whole-brain volumes were collected. In addition, a high-resolution T1 MR sequence was collected with the following parameters: TR=2.3 s, TE=2.98 ms, IT=900ms, matrix size 256×256, voxel size : 1×1×1 mm, 160 slices, Bandwidth 240Hz/voxel.

#### Audio-tactile speech comprehension Training

The training session took place in a room at ENU after the SRT estimation tests and after the first task-fMRI run was completed. The task in the training was to repeat sentences, from the database of 148 vocoded HINT sentences presented against background speech noise (some sentences were new and some were the same as those used already in the experiment). Each sentence played in the headphones was accompanied with congruent low-frequency vibrotactile inputs delivered on two fingertips of the right hand via the tactile device (same device as the one used in the MR scanner). The SNR during training corresponded to the individual SRT value, as established before the training for the ATc condition for each person. During the session each sentence was presented up to 2 times in a row and if the person was unable to repeat it correctly, the sentence was presented as text on the PC screen in front of the participant (black font, white background, middle of the screen). Then the person listened to the sentence again, while also looking at it on the screen. Each sentence requiring such feedback was stored in the database and replayed at the end of the session. The session continued until all 148 sentences were repeated correctly without feedback. For all subjects, the training was completed within 30–45 min. The feedback was determined to be visual as opposed to auditory, as in the future the authors wish to apply the same, albeit adjusted, study procedures in a group of hearing impaired participants.

### Data analysis

#### Behavioral data analysis

The SRT values obtained by the participants in the behavioral speech comprehension tests before and after the training session were compared between task conditions within the Pre or Post session (A, ATc, ATnc; Wilcoxon ranked-sum tests), and between sessions (A Pre vs A Post, ATc Pre vs ATc Post, ATnc Pre vs ATnc Post; Wilcoxon signed-rank tests).

#### MRI data analysis

First, in order to elucidate the brain network involved in speech comprehension tasks, fMRI data were analyzed using SPM12 using an univariate approach (https://www.fil.ion.ucl.ac.uk/spm/software/spm12/). The anatomical images were scalp-striped, corrected for field inhomogeneities and transformed into MNI space. For functional runs the following analysis steps were applied: slice time correction, removal of linear trends, and 3D motion correction to the first volume of each run using rigid-body transformation. Each run was coregistered with the native-space anatomical image (T1) of the participant, and subsequently warped into MNI space with the same affine+non-linear transformation used for warping anatomical data. To create group-level activation maps, the MNI-warped functional images were first smoothed with a 3D Gaussian kernel of 5 mm FWHM and then entered into a hierarchical random effects analysis (RFX GLM; Friston et al., 1999). Each experimental condition’s predictor was modeled by convolving the boxcar predictor describing the condition’s time-course with a standard hemodynamic response function (HRF). In addition, the model included motion-related effects (using the z-transformed motion estimates generated during preprocessing). Activation maps (one-sample t-maps) for each run separately were thresholded at p=0.001 (voxel) and FWE=0.05 (cluster). The following contrasts were applied next for between-condition and between-session comparisons: A PRE vs ATc PRE, A PRE vs ATnc PRE, ATc PRE vs ATnc PRE, A POST vs ATc POST, A POST vs ATnc POST, ATc POST vs ATnc POST, A PRE vs A POST, ATc PRE vs ATc POST, ATnc PRE vs ATnc POST, ATc vs ATnc vs A PRE, ATc vs ATnc vs A POST.

Second, in order to evaluate functional connectivity patterns between brain regions, CONN 22a (Connectivity Toolbox, https://web.conn-toolbox.org/) was used for data analysis. Preprocessing of the functional data included: slice timing correction, motion correction, scrubbing, linear detrending, band-pass filtering (0.008 Hz < f < 0.09 Hz), co-registration to individual T1 structural scans, spatial normalization (volumetric) to a standard brain space (MNI template), and spatial smoothing (6 mm Gaussian kernel). Denoising was applied with default settings. A group-ICA analysis was performed using 40 components and the G1 FastICA + GICA3 back-projection procedure with 64 dimensionality reduction. Next, independent components were selected that represented certain brain networks and regions (e.g. : auditory, visual, sensorimotor, language) using the “compute spatial match to template” tool and based on visual inspection. Functional connectivity maps were compared between sessions and between runs (A PRE vs ATc PRE, A PRE vs ATnc PRE, ATc PRE vs ATnc PRE, A POST vs ATc POST, A POST vs ATnc POST, ATc POST vs ATnc POST, A PRE vs A POST, ATc PRE vs ATc POST, ATnc PRE vs ATnc POST). The subsequent statistical analysis used permutation with the following parameters : voxel-threshold p <0.01, cluster-mass p-FDR corrected p < 0.05.

For all the results the regions were labeled using the Harvard-Oxford Brain Atlas or the AAL atlas.

## Results

### Results of the speech comprehension tasks : lowest SRT values in the congruent audio-tactile task after training

Before training the SRT values were highest (indicating poorest performance) for the A Pre test condition (M=17.5 dB, SD=7.49 dB), as compared to ATc Pre (M=11.1 dB, SD=4.4 dB) (z=2.67, p=0.008) and to ATnc Pre (M=10.7 dB, SD=5.7 dB) (z=3.34, p=0.001), with the latter two not different from one another (p>0.05). After training the SRT was lowest (indicating best performance) for the ATc Post condition (M=2.6 dB, SD=6.3 dB), as compared to the ATnc Post condition (M=7.7 dB, SD=4.9 dB) (z=2.742, p=0.006) and the A Post condition (M=7.5 dB, SD=5.3 dB) (z=2.741, p=0.006). SRT values for A Post and ATnc Post were not statistically significantly different from one another (p>0.05). SRT values significantly improved for the A condition (z=3.48, p<0.001) and for the ATc condition (z=3.48, p<0.001). There was no significant improvement in performance in the ATnc task (p>0.05).

### Results of the fMRI data analysis

We present below functional brain networks involved in the speech comprehension task and multisensory integration revealed by univariate GLM analysis, and changes in functional connectivity related to the task condition and training.

### fMRI univariate functional analysis reveals a typical speech network in all test conditions

One-sample t-test results of six fMRI runs during which the participants were repeating sentences were depicted in Figure 2. Common regions revealed for all speech comprehension conditions included bilaterally in the cortex: superior temporal lobes (STG, pSTS, Heschl gyri, TP), superior marginal gyri (SMG), middle temporal gyri (MTG), inferior frontal gyri (IFG, opercular and triangular), precentral gyri (area 4), postcentral gyri (area 3b, 3a), Rolandic operculi (OP1-4), as well as the middle cingulate gyrus, and the supplementary motor area (SMA); in the cerebellum bilaterally : vermis 4-8, cerebellum lobes 4-8, crus 1-2. For the A PRE test and A POST test conditions there was additional involvement of the visual calcarine and the lingual gyri (V1, V2, V6), and subcortically in the thalamus, palladium, putamen, caudate, anterior insula, superior and inferior colliculus and cochlear nuclei. In the A POST test, there was no/less activation detected in the basal ganglia and subparts of the cerebellum (eg. left crus 1-2, vermis 4-8, right 4-5 lobe), as opposed to A PRE. For ATc PRE, the revealed network was very similar to A PRE but there was no activation found in the visual system and several divisions of the cerebellum (e.g. the vermis). After training (ATc POST), the responses in the calcarine, as well as in all basal ganglia and the cerebellum were very limited (e.g. no activation in the left crus 2, lobes 4-5 and 7, vermis). Otherwise, for the congruent audio-tactile task, the activations demonstrated a very similar pattern before and after training. During the ATnc condition, the revealed network was very similar to A PRE and ATc PRe, but the size of activation in most regions, including IFG, SMG, insula, MTG, Pre and Post-Central gyri was much larger. After training (ATnc POST), the found regions were almost identical as in ATnc PRE. In all conditions, the size of activation after training diminished. A table with details regarding the revealed activation maps (e.g. MNI coordinates, t-values, etc.) is available upon request from the corresponding author. No statistically significant differences were found when the results of the 6 fMRI runs were compared with one another (two-sample t-tests, cluster-wise p<0.05 FWE-correction for multiple comparisons).

**Figure 2.**
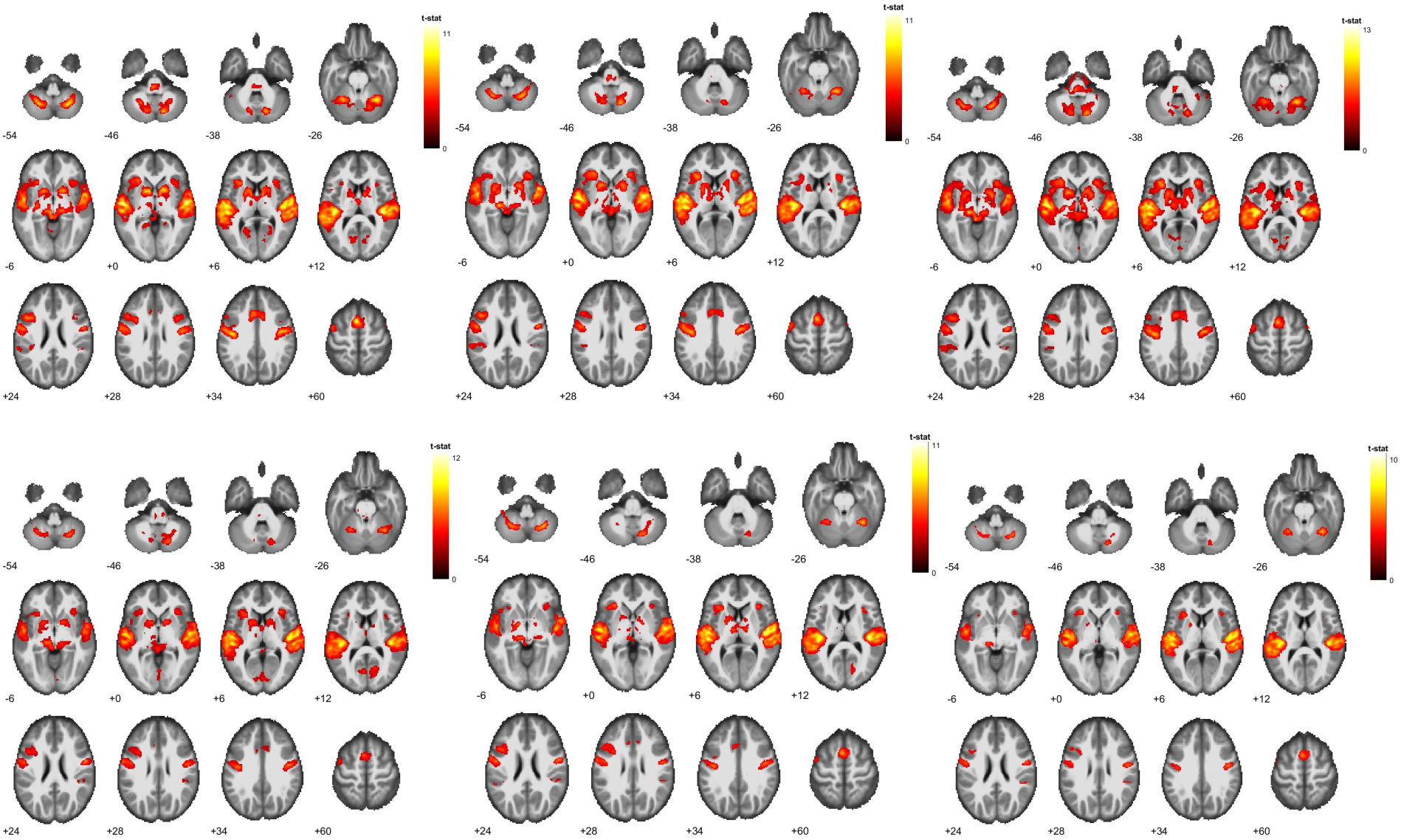
Results for three fMRI speech comprehension tasks; first row : Audio PRE, Audio-Tactile PRE congruent (ATc), Audio-Tactile PRE non-congruent (ATnc), second row : Audio POST, Audio-Tactile POST congruent (ATc), Audio-Tactile POST non-congruent (ATnc); GLM univariate analysis; p<0.001, cluster FWE <0.05.

### fMRI functional connectivity results show an effect of training and an effect of the applied test condition

#### Effect of training (pre vs post) : FC changes in the visual system

After training (in comparison to before), FC was *increased* for the congruent audio-tactile test condition (ATc) between the region composed of (mainly) bilateral iLOC/sLOC, lingual gyrus (LG), occipital fusiform gyri (OFG), and right middle & inferior temporal gyrus (posterior MTG, ITG), as well as left OP [seed], and an area encompassing parts of the right inferior/superior lateral occipital cortex (i/sLOC) and right occipital pole (OP) [target]. After training functional connectivity was *decreased* for ATc between bilateral OP, intra-calcarine cortex (ICC), LG, and left i/sLOC [seed] and the same target in the right-hemisphere occipital area; as well as between an area composed of parts of bilateral OP, iLOC/ sLOC, LG, ICC, cuneus [seed] and left i/sLOC [target]. No difference in FC was found as an effect of training for the A and the ATnc speech test conditions. See Figure 3.

**Figure 3.**
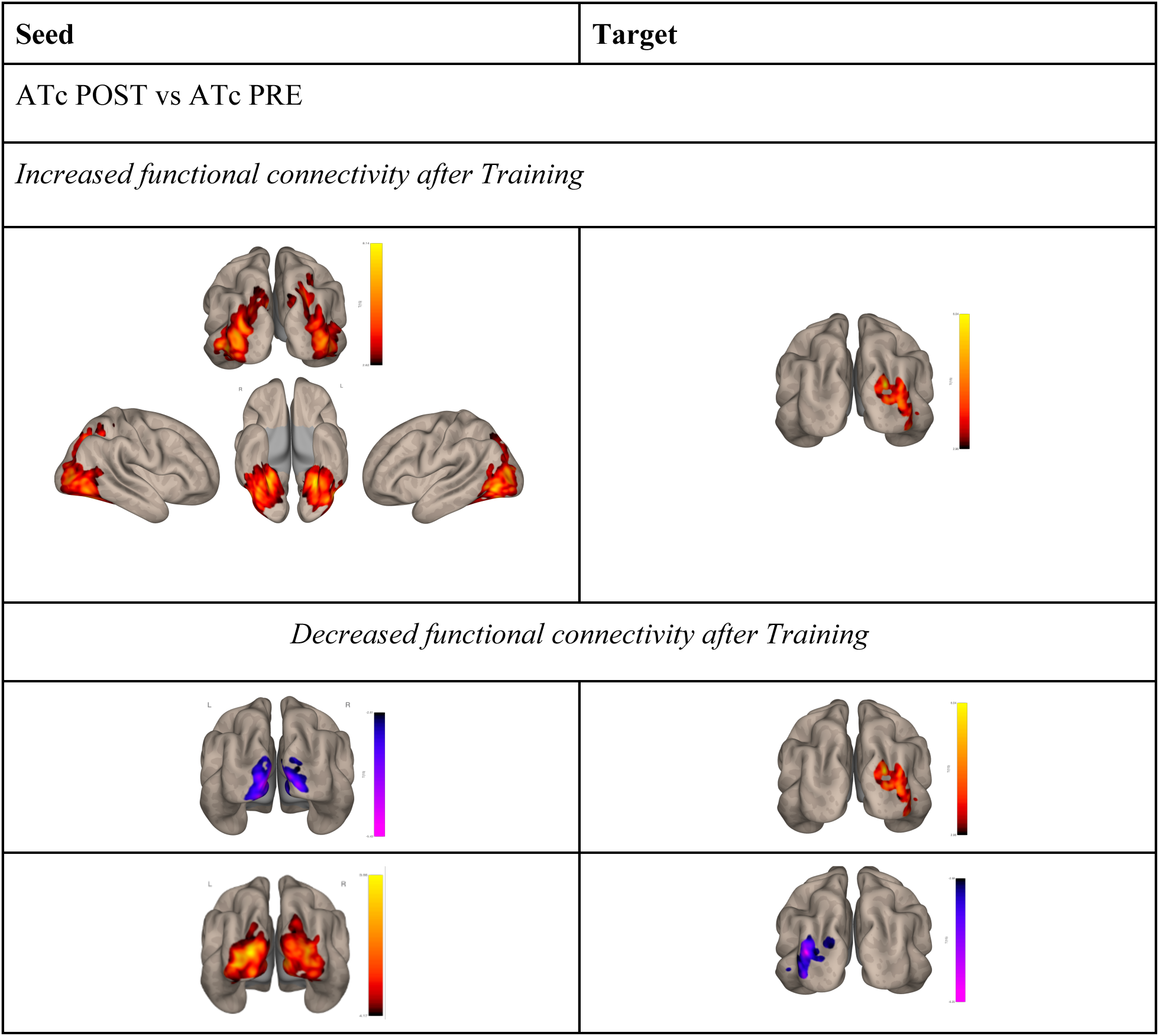
Results of the functional connectivity analysis of task-fMRI data for speech comprehension: effect of training for the congruent audio-tactile test condition; if both seed and target are the same color (red or blue), this indicates increased FC; if they are different colors (blue-red or red-blue), this indicates decreased FC.

#### Effect of condition (between conditions): FC changes in the sensorimotor, visual, and high-order frontal brain regions

Before training FC was *increased* for the multisensory ATc speech test (as compared to audio only) between a network composed of parts of of bilateral precentral & postcentral gyri (PreCG & PostCG, respectively), Supplementary Motor Area (SMA), Superior Frontal Gyrus (SFG), Superior Parietal Lobule (SPL), and Parietal Operculum (Op) (further as SM for sensori-motor network, [seed]) and a network combining bilaterally sLOC, SPL, Angular Gyrus (AG), posterior Superior Marginal Gyrus (pSMG), LG & cuneus, ICC, Superior Calcarine Cortex (SCC), as well as right iLOC and precuneus [target]. Before training *increased* FC was also shown for combined ATnc (as compared to audio) between the SM network [seed] and bilaterally sLOC, SPL, Angular Gyrus (AG), and precuneus (same parts as above) [target]. After training, there was *increased* FC when ATc was compared to A between the SM network [seed] and a network consisting of parts of bilateral frontal pole (FP), SFG (different part than earlier), frontal medial/orbital cortex, bilateral cingulate cortex, precuneus, and on the left side : anterior MTG, AG & sLOC (same parts as above) [target]. No difference between ATc and ATnc, or between ATnc and A FC networks was found either before or after training. See Figure 4.

**Figure 4.**
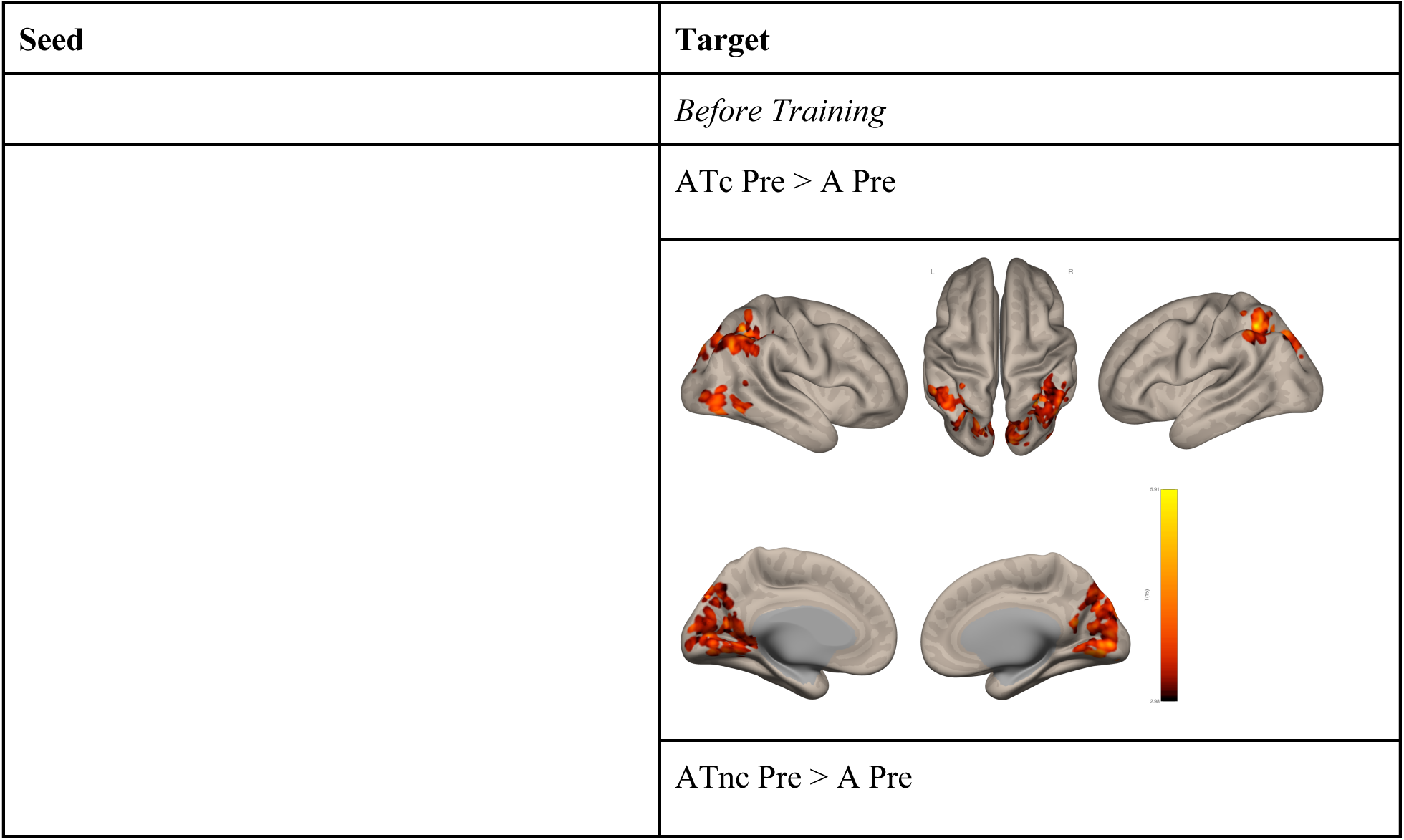
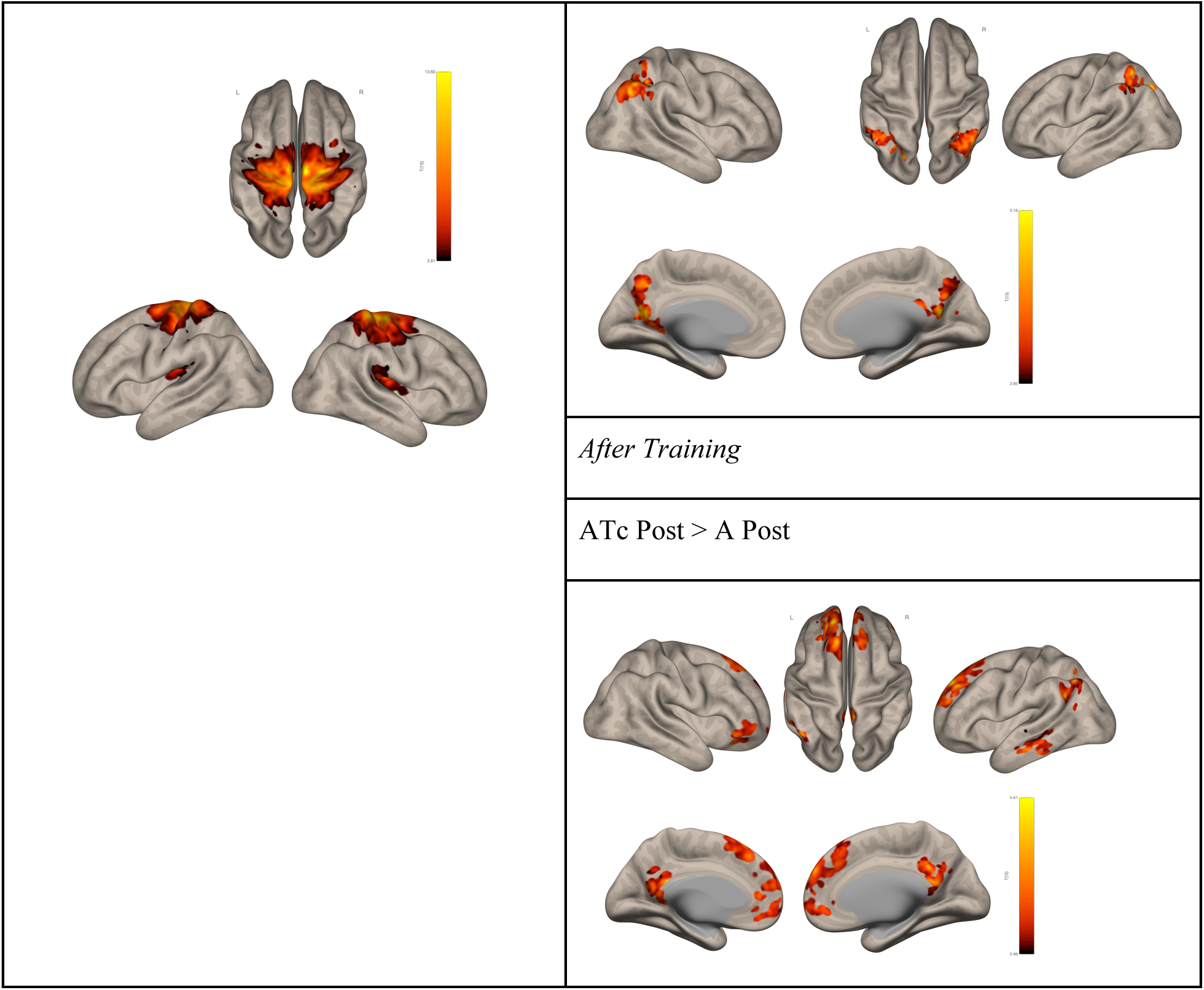
Results of the functional connectivity analysis of task-fMRI data for speech comprehension: effect of the test condition before and after training (congruent audio-tactile vs audio only); *if both seed and target are the same color (red or blue), this indicates increased FC; if they are different colors (blue-red or red-blue) this indicates decreased FC*.

## Discussion

### Speech comprehension improvement after audio-tactile training

As hypothesized, the behavioral outcomes of the study show that typically hearing individuals were able to recognize vocoded speech in noise, both on its own and when accompanied with vibrotactile speech input, and improved in comprehension after a dedicated training session. The training-related gain, reflected in decreased SRT values, was significant for the auditory test condition and when speech was delivered as congruent audio-tactile stimulation, whereas the scores in the control condition (with speech signals paired with non-congruent vibrations) remained largely unchanged. Even though the training session involved multisensory inputs, the auditory component was always present and therefore practiced by the participants, leading to improved understanding through hearing only, as well. Indeed a number of studies have shown that intelligibility of distorted speech or speech in noise can improve through experience and training in various contexts (e.g. Casserly and Pisoni, 2015, Erb et al., 2013; Guediche et al. 2014). Before audio-tactile speech was trained, comprehension was found to be at a similar level regardless whether the additional tactile information was congruent or non-congruent with the auditory speech signal, and for both these conditions better than for the auditory speech alone. After the training the congruency significantly improved the scores, and the lowest mean SRT values, indicating highest performance in the most challenging acoustic conditions, were obtained for the congruent audio-tactile test condition. These results together suggest that: a) training led to increased discrimination between the useful congruent vibrotactile stimuli and the less useful or distracting non-congruent tactile stimuli, b) the congruent auditory and vibrotactile speech inputs were successfully integrated, leading to improved speech comprehension in noise. Combining both sensory inputs efficiently required practice, especially that the tactile inputs representing speech were an utterly novel experience to the participants. In our previous work with the same set-up we showed that such multisensory benefit also occurs without any training, but is less profound (Ciesla et al. 2019). We also previously showed that multisensory audio-tactile training as compared to unisensory auditory training leads to improved speech comprehension much more efficiently (Ciesla et al. 2022). The latter effect is in line with other studies showing superiority of multisensory (traditionally, audio-visual) over unisensory speech training (e.g. Bernstein et al. 2022; Lidestamm et al. 2014). We believe that integration of the audio-tactile stimuli facilitated speech understanding in two ways; first by complementing the distorted speech signal with low-frequency temporal information (envelope) available in the vibrotactile input, and second by improving the contrast between the target sentence and the background noise (Crommett et al. 2019). The latter mechanism might also be the reason for improved comprehension (lower SRT) before training for the non-congruent audio-tactile test condition, as compared to unisensory auditory speech. Also in the “typical” multisensory context of speech perception, auditory comprehension has often been shown to improve when accompanied with congruent visual inputs (i.e. looking at the face and/or the body of the speaker), and especially in the context of reduced intelligibility of the auditory signal, convergent with the rule of inverse effectiveness of multisensory integration (Sumby and Pollack 1954; Hickock et al. 2018; Meredith and Stein 1986, Lidestamm et al. 2014, Bernstein et al. 2013). The presented tactile intervention can potentially provide the same type of benefit to the hearing impaired, with possibly more rapid effects, as the tactile inputs contain exactly the same type information (a temporal signal fluctuating in amplitude and intensity) as the auditory inputs (Huang et al. 2017).

### Typical speech networks revealed with univariate fMRI analysis

All speech comprehension tasks, both auditory and audio-tactile, before and after training, produced similar patterns of activation, with no statistically significant differences between them. The shared revealed networks included the classical left-hemisphere dorsal articulatory-phonological pathways (connecting the posterior STG to the Broca area in the inferior frontal cortex, and pMTG/STG/SMG to the premotor cortex), as well as the ventral semantic pathways (from the inferior frontal cortex to STG/MTG, and to anterior STG/TP), all converging in the auditory cortex in the superior temporal lobe (Friederici 2013, Hertrich et al. 2020; Hurley et al 2015). Additional revealed regions included the right IFG subserving semantic/pragmatic monitoring (Neef et al. 2016), SMA and pre-SMA engaged in cognitive and low-level motor control in speech production (Hertrich et al. 2020), as well as bilateral SMG participating in phonological analysis, articulation and multisensory integration (Oron et al. 2016; Oberhuber et al. 2016). The revealed subcortical structures, cerebellum, the insula, and the basal ganglia have often been shown as involved in language processing, but also in general learning (Hertrich et al. 2020; Price et al. 2012). The visual cortex recruitment in speech perception, found here for auditory conditions, is discussed in further paragraphs. We speculate that the lack of difference in activation between the unisensory and multisensory speech conditions might be partially related to the shared neural mechanisms of encoding both auditory, tactile and audio-tactile inputs in the auditory cortex or the posterior temporal lobe. Several classic studies suggested such an effect for high-frequency non-speech vibrotactile stimuli in both healthy and deaf individuals (e.g. Caetano and Jousmäki, 2006; Foxe et al. 2009, King et al. 2019, Kayser et al. 2005; and see also a similar finding for trained tactile speech in Damera et al. 2023). A similar suggestion has also been put forward for audio-visual speech integration in pSTS (Hertrich et al. 2020; Beer et al. 2013). This however needs to be investigated further with a dedicated paradigm including a “tactile only” condition. At the same time, the training might have been too short to induce changes in levels/size of activations across brain regions. This, specifically that group-level functional MR results are highly sensitive to inter-subject variability in activation loci, size, and amplitude, which can obscure focal functional effects and between-group or between-task differences (Mankin and Krakauer 2023; Nath et al. 2012, Beauchamp et al. 2004).

### Training-related functional connectivity changes in the “visual system” for the trained audio-tactile task

We show that training-induced functional connectivity (FC) changes were (only) found for the trained ATc speech condition, and specifically between areas within the “visual system”. Decreased FC was shown within early visual regions (V1-V3), whereas FC between V2/V3 and subcomponents of the dorsal (intraparietal sulcus/ IPS, MT, Extrastriate Body Area /EBA) and ventral (the fusiform gyrus, V4, iLOC) visual streams was increased (Goodale et al. 1992, Sheth et al. 2014). This, even though no language-related visual input was present during the fMRI experiment. Although traditionally V1-V3 are considered low-level unisensory areas, they have been shown engaged in simple auditory and tactile tasks, indicating their multisensory or amodal potential (e.g. Burton et al. 2004; Murray et al. 2016; Landelle et al. 2023; Martuzzi et al. 2007). Furthermore, early visual regions were found to be involved in the processing of spoken language in the congenitally blind population (Amedi et al. 2003; Bedny et al. 2015; Roeder et al. 2002; Vetter et al. 2014; Beck et al. 2023; Burton et al. 2002), with the amount of activation proportional to the syntactic difficulty/semantic content, or the blind individual’s verbal-memory skills (Roeder et al. 2002 and Amedi et al. 2003, respectively), as well as in tactile language processing, such as when reading Braille (e.g. Sadato et al. 1996; Reich et al. 2011). There are also several studies showing processing of non-visual speech information in early visual areas in the sighted population, indicating existence of such interactions in a healthy brain as well (potentially unmasked by sensory deprivation; Makin and Krakauer 2023; Bedny et al. 2011; Seydell-Greenwald et al. 2023; Merabet et al. 2008). As an example, early visual recruitment was found in the sighted for Braille reading following blindfolding for 5 days (Marabet, 2008; Pascual-Leone and Hamilton, 2001), as well as for auditory speech perception (Seydell-Greenwald et al. 2023). In addition, a series of studies showed that auditory sounds, including speech, can be decoded (categorized) based on activity in the early visual cortex (V1-V3) in blindfolded sighted individuals and in the blind (Vetter et al. 2014, Vetter et al. 2020). One possible mechanism of the audio-visual interactions might be that sound stimulation and the associated images generate feedback to the early visual cortex, carrying abstract and possibly semantic information. This might specifically be the case of audio-visual speech processing, i.e. with concurrent lip-reading. In everyday life, especially in challenging acoustic situations, the alternative sensory information available for speech understanding is typically visual. In the current study, after training, the network with increased FC included V2, MT, EBA (LOC) and the fusiform gyrus, all previously shown for speech perception, i.e. when encoding high-resolution details of moving lips, body gestures and facial expressions/language-related cues, respectively (Bernstein et al. 2022, Zhang et al. 2016, Kitada et al. 2014; Hickok et al. 2018, Van der Stoep et al. 2020). It has been speculated that the linguistic information may reach these extrastriate visual regions from language regions in the temporal lobe, through prefrontal cortex and/or language-relevant nuclei of the thalamus (Bedny et al. 2011; Liu et al. 2007). Taken altogether, it is possible that when exposed to the novel audio-tactile speech input, the dorsal and ventral visual streams both become engaged automatically due the life-long audio-visual associations established for speech perception. This assumption is even more reasonable, with the task being acoustically challenging (involving distinguishing the target speech signal, by itself distorted, from competing background noise), and thus encouraging the use of alternative sensory information and increasing the chance for multisensory integration. At the same time, the internal connectivity of V1/V2 decreased after training, possibly due to the actual lack of any visual inputs during task performance (as opposed to tactile). The fact that early visual regions were only revealed for the auditory speech tasks in the classical univariate analysis provides further support to this speculation.

#### Functional connectivity changes related to audio-tactile integration involve multiple brain regions

The second of the revealed effect pertains to the increased FC of the sensorimotor network (SM), and specifically both the primary and secondary regions representing the trunk, legs and possibly the fingers, during audio-tactile integration (audio-tactile condition vs audio only), with several other brain areas. Before training, for both ATc and ATnc, the connectivity of the SM network was increased with a localized bilateral area in posterior parietal cortex (PPC), a subregion known for its multiple functions. First of all, part of the detected hub can be considered the association somatosensory cortex, consistently found to engage in higher-level tactile operations, including Braille reading, shape recognition, roughness estimation, tactile attention, etc. (Kim et al 2014, Bodegård et al., 2001; Li Hegner et al., 2010, Burton et al. 2004). Indeed, in our experiment, the newly experienced tactile signal must be carefully examined for the temporal or speech features it represents. The PPC (specifically regions such as the Parietal Reaching Region and the Parietal Eye Field, adjacent to the Intraparietal Sulcus) is furthermore believed to engage in computing motor commands for the body, such as reaching/grasping or directing visual attention through eye movement, based on the arriving multisensory inputs (e.g. Orban et al. 2021, Azañón et al. 2010). The IPS in general (also revealed to have increased connectivity with V2 following training in the current study; see previous paragraph), is part of the dorsal visual stream responsible for spatial vision i.e. locating objects and one’s body position in space, or in a more general sense, amodal spatial attention (Sheth et al. 2014; Goodale and Milner 1992). Beyond the parietal hub, before training for the ATc speech task, the connectivity was also enhanced between the SM network and medial early visual areas (V1-V3), bilaterally. While no explicit visual input was given during the task in the scanner, direct connections have been demonstrated from auditory core and parabelt to V1 (Falchier et al., 2002) and from V2 to the caudal auditory cortex in monkeys (Falchier et al., 2010). These connections can represent, among others, a feedback mechanism in the dorsal visual pathway, representing multisensory extra-personal space and motion, and possibly exist (among other functions) to enhance general spatial orientation (Azevedo et al., 2015; Cate et al., 2009). With respect to the revealed multisensory areas specifically, we found both bilateral SMG/AG as well as the right-hemisphere iLOC to have increased connectivity with the SM map before training (right AG was also found to have enhanced FC with the SM hub before training for the non-congruent audio-tactile condition). SMG and AG are regions often found to engage in phonological and semantic language operations, respectively, but also other high-level functions (Oberhuber et al. 2016, Seghier et al 2013), whereas iLOC engages in multisensory (albeit mainly visuo-tactile) tasks including grasping and visually guided finger movement (e.g. Amedi et al., 2002; Kassuba et al., 2013; Jao et al. 2015; Brand et al. 2020), but also when perceiving passive vibrotactile stimulation on the hand (possibly related to object recognition and manipulation; Tal et al. 2016). The nearby V4 region, often reported for texture perception, might reflect the similarity between texture and active exploration of vibrotactile inputs on the fingertips, both employing the same mechanoreceptors (Landelle et al. 2023; Bensmaia et al. 2003). While for both the congruent and non-congruent multisensory conditions the FC was increased between SM and PPC/SMG/AG, only for the former also early visual regions and iLOC were engaged. This might again suggest a specific role of the visual system in successful multisensory integration (Murray et al. 2016). All these results together may represent a potential neuronal mechanism accompanying the audio-tactile speech comprehension task before a dedicated training session. On the one hand, it seems that in order to perform the novel task the participants need to actively localize all their body parts in space, including the palms and the fingertips on the sides of their body (a process similar to reaching or grasping). In parallel or subsequently, they engage in the analysis of the novel tactile inputs, and try to combine these with the auditory inputs received by the ears, in multisensory brain regions. After training, i.e. after the vibrotactile language input is already more familiar and practiced, the mechanism of audio-tactile integration becomes more specifically language-oriented, albeit still engaging the SM system. Specifically, the same sensorimotor network as before training shows enhanced connectivity with the left temporal-parietal junction, left posterior middle temporal gyrus (pMTG), as well as several bilateral subregions of the frontal cortex (dorsomedial prefrontal cortex, DLPFC and orbitofrontal cortex, OFC). While the DLPFC and PPC, through multiple direct connections, comprise the fronto-parietal/central executive network, serving, among others, sustained attention, executive functions, working memory (e.g. Menon et al. 2021), DLPFC can also engage in language tasks specifically (e.g. in sentence vs single word comprehension), through numerous connections with both dorsal and ventral auditory pathways (Hertrich et al. 2020). OFC has been recently shown to both respond to sound itself as well as actively modify processing in the primary auditory cortex in rodents, through dedicated top-down connections (e.g. Mittelstadt 2023), with possibly similar functions in humans. In addition, OFC was also found to be one of neuronal correlates of listening effort (Rosemann & Thiel 2020; Wikman et al. 2022), especially relevant in the task in the current study. Also the revealed left TPJ, on the junction of pSTG/AG/SMG/IPL, is believed to support recognition of degraded or ambiguous speech, in the process of perceptual learning (Davis 2016, Jurewicz et. al 2020). Posterior MTG is a typical language area involved in semantic retrieval, control and lexical decision making, densely connected to other language areas such as the inferior frontal gyrus, premotor cortices, as well as SMG and AG (e.g. Xu 2015).

## Summary and conclusions

In summary, we showed that complementing distorted auditory signals with speech-extracted vibrotactile inputs improved speech comprehension in noise, and specifically after a dedicated training session. The improvement was accompanied with changes in functional connectivity, but no effects were revealed in brain activation patterns. The demonstrated effects with respect to functional connectivity were partially in line with our hypotheses. As expected, we showed a substantial overlap between the set of regions engaged in audio-tactile speech processing and the established audio-visual speech network, including vast involvement of the dorsal and ventral “visual” stream (V1/2, MT, EBA, iLOC). At the same time, however, this network was still found involved in the audio-tactile task after training. Second, against expectations, the applied analysis did not show any change in FC between sensorimotor areas and the auditory cortex or pSTS specifically. We speculate that in order to both weaken the connectivity within the visual system and strengthen the direct audio-sensory pathway of communication, a longer training regime of the novel multisensory context of speech is required. For audio-tactile integration specifically, FC analysis, as hypothesized, revealed both parietal and frontal integration and language-related areas, but also an unexpectedly sizable area representing the whole body map (even though this cluster was an outcome of ICA analysis and therefore does not represent an activation area specific for task performance). As discussed, the involvement of this hub along parietal regions, was possibly related to the body and space perception mechanism required to combine the two incoming signals, auditory and tactile. In conclusion, the results of the study demonstrate that an adult brain maintains the capability to combine newly available sensory information (here tactile) with the existing and long-trained sensory modality (here auditory) for a novel task (Heimler and Amedi 2020). The process is relatively rapid, possibly due to the number of similarities between the two signals, both representing the same temporally changing input, thus opening new rehabilitation opportunities for the hearing impaired (Crommett et al. 2019. Crosse et al, 2016). In addition, brain plasticity related to perceptual learning is shown to be mainly built upon an existing structural and functional blueprint for connectivity (that can nevertheless change through reweighting, strengthening and weakening; Makin and Krakauer 2023).

## Limitations of the study

In all participants the auditory speech comprehension task always preceded the two audio-tactile tasks. This was done in order to first familiarize the participants with the speech material, and then to keep the procedure consistent. This might have contributed to the lower results (i.e. higher mean group SRT values) being obtained in the first auditory task before training. However, we believe that the effect was minimal, and also not relevant after training, i.e. following exposure to more than 200 sentences, including during the training session focused on multisensory integration.

## Acknowledgements

This research was supported by Polish Ministry of Science and Higher Education MOBILNOŚĆ PLUS V grant (1642/MOB/V/2017/0) to K.C., an ERC Consolidator Grant (773121 NovelExperiSense) and a Horizon GuestXR (101017884) grant to AA.

## Authors contributions

KC : writing-original, writing - review & editing; conceptualization; data curation; formal analysis; investigation; methodology; project administration; software; supervision; visualization

TW : writing - review & editing; conceptualization; formal analysis; methodology; software; visualization

AA: writing - review & editing; funding acquisition; methodology; resources; supervision

## Competing interests

The authors declare no competing interests.

## Notes

### Competing Interest Statement

The authors have declared no competing interest.

